# From patterning to secretion: Kv2.1 subunits as regulators of zebrafish hatching gland morphogenesis and function

**DOI:** 10.64898/2025.12.18.695154

**Authors:** Ruchi P Jain, Rosa R Amini, Vladimir Korzh

**Affiliations:** Laboratory of Neurodegeneration, International Institute of Molecular and Cell Biology in Warsaw, Poland

**Keywords:** zebrafish, hatching gland, *kcnb1* and *kcng4b*, exocytosis, secretion, abnormal patterning, morphogenesis

## Abstract

Zebrafish hatching, a critical developmental milestone, occurs around 48-72 hours post-fertilization (hpf). It is regulated by the specialized secretory organ called the hatching gland. Voltage-gated potassium channels (Kv) are known for their roles in maintaining plasma membrane potential and regulating intracellular protein traffic and secretion. Previous studies on zebrafish mutants of Kv2.1 channel subunits - the electrically active α subunit Kcnb1 and the modulatory subunit Kcng4b - revealed antagonistic functions in the development of brain ventricles, ear, and Reissner fiber. In this study, we investigated their functional role in the hatching gland. The loss of either subunit resulted in a significant delay in normal hatching. Using *in situ* hybridization and immunohistochemistry, we show that both mutants exhibited severe defects in the hatching gland patterning, including a reduced number of hatching gland cells. The mutants displayed changes in the transcript levels of several hatching gland markers and reduced cell proliferation in this organ. These developmental defects were intensified by a late-stage functional failure characterized by decreased cathepsin synthesis, reduced proteolytic activity, and delay in the period of secretion in both mutants. Together, our findings establish that Kv2.1 subunits, Kcnb1, and Kcng4b are essential during the development of the zebrafish hatching gland and its secretion.

## 1 INTRODUCTION

Hatching is a pivotal step in the development of oviparous animals, marking the transition from a protected embryonic to a free-swimming larval stage. In zebrafish, this process typically occurs between 48 and 72 hours post-fertilization (hpf), when the larvae escape the protective envelope, known as the chorion (1,2). A functionally analogous event, termed blastocyst hatching, occurs in viviparous species (e.g., human), where the embryo hatches out of the zona pellucida around 5-7 days, a process essential for successful implantation (3,4). Failure of this essential transition often results in delayed development or developmental arrest, highlighting hatching as a sensitive and vital checkpoint of normal embryonic progression (5,6).

Hatching is a process that exhibits significant plasticity and is influenced by a variety of internal and external factors. These factors can be broadly classified as: (i) environmental (temperature, oxygen, humidity, and nutrition); (ii) mechanical (embryo’s pressure on the chorion); and (iii) chemical (specialized hatching enzymes to degrade the chorion). The latter is represented by a cocktail of specialized proteolytic enzymes (4,7,8). In zebrafish, these enzymes are synthesized and released by a transient embryonic organ known as the hatching gland (HG). This specialized tissue develops from the anterior-most portion of the midline mesendoderm (polster), a lineage that also gives rise to the prechordal plate, floor plate, and notochord (9–11). During embryogenesis, the HG cells undergo coordinated migration and differentiation to form a cohesive tissue band on the enveloping layer. These cells function as an endocrine secretory unit (12). They are characterized by the presence of large secretory granules, which store synthesized proteases, including cathepsin LB (Ctslb) and hatching enzyme 1, tandem duplicate 1 (He1.1) (13–15). The function of the HG is the unique mode of terminal enzyme release known as holocrine secretion, in which the cell ruptures and dies upon secreting its contents (16,17). Given that this dramatic and irreversible event is triggered by a complex cascade of regulatory signals (2,7), the HG serves as a highly sensitive model for studying the downstream roles of membrane potential and ion channels in regulating specialized cellular function and precise developmental timing.

Voltage-gated potassium (Kv) channels are essential membrane proteins that regulate cellular excitability and function by controlling the flow of K^+^ ions across the plasma membrane (18,19). These channels are crucial for maintaining the cell’s resting potential as well as cell proliferation, migration, and apoptosis (20). The Kv2.1 channel is a significant component of the delayed rectifier current (21). KCNB1, which encodes the electrically active subunit of the Kv2.1 channel, is linked to Developmental Epileptic Encephalopathy (DEE). There are more than 50 heterozygous pathogenic *de novo* variants identified, which cause a wide range of symptoms, including seizures, autism, and developmental delay in patients (22–24). Beyond its primary electrical role, Kcnb1 forms specialized clusters at the plasma membrane essential for regulating the targeting and release of secretory granules during exocytosis (25). KCNB1 forms heterotetramers with electrically inactive (or modulatory) subunits, such as Kv6.4 (encoded by *KCNG4*). To form functional tetramers, KCNB1 acts as a shuttle that traffics the silent subunit KCNG4 from the endoplasmic reticulum (ER) to the plasma membrane. The co-assembly of KCNB1 and KCNG4 modulates channel current density (26–29). Deletion of *Kcng4* in mice affects spermiogenesis (30). Zebrafish possess two paralogs of *Kcng4*: *kcng4a* and *kcng4b* (31). During zebrafish development, *kcnb1* and *kcng4b* exhibit overlapping expression patterns in various tissues. Kcnb1 and Kcng4b play an antagonistic role in the development of brain ventricles, ears, and Reissner fiber (32–35). Crucially, although one transcriptomic atlas suggests an expression of these genes in the HG (36), this finding is disputed by others (37), leaving the functional roles of Kcnb1 and Kcng4b in HG development and secretion unexplored.

In this study, we utilized two zebrafish mutants: *kcnb1^-/-^*represents the loss-of-function of the electrically active α-subunit Kcnb1, and *kcng4b^-/-^* represents the loss-of-function of the silent subunit (Kcng4b). Using these tools, we addressed the functional role of Kv2.1 in HG. We demonstrate that loss of either Kv2.1 subunit disrupts the core processes of HG development, including patterning, cell proliferation, and synthesis of proteolytic enzymes, and ultimately compromises the terminal holocrine secretory mechanism, leading to a hatching delay.

## 2 MATERIAL AND METHODS

### 2.1 Animals

Zebrafish (*Danio rerio*) lines were maintained at 28°C, on a 14:10 h light/dark cycle in E3 medium with 0.0015% methylene blue, following established standard husbandry protocols (38) in the Zebrafish Core Facility of the International Institute of Molecular and Cell Biology, Warsaw (license No. PL14656251). All procedures involving embryos and larvae adhered to the guidelines of the Polish Laboratory Animal Science Association and European Communities Council Directive (63/2010/EEC). Three groups of zebrafish were used: wild-type (WT), the *kcng4b* mutant *kcng4b^waw304^* (39), and the *kcnb1* null mutant *kcnb1^sq301^* (32). We only used the homozygous mutants (*kcnb1^-/-^* and *kcng4b^-/-^*). For manuscript readability, these mutants will be referred to using their gene shorthand, *kcnb1* and *kcng4b*, respectively.

For all experiments, embryos were dechorionated with pronase (#11459643001, Roche) at a final concentration of 1 mg/mL, unless otherwise specified. For all imaging purposes, embryos were treated with 0.2 mM phenylthiourea (PTU, #L06690, Thermo Fisher) starting at 8 hpf to block pigmentation.

### 2.2 Hatching assessment

Zebrafish embryos were incubated in E3 medium at 28°C, and the medium was replaced every 24 hours. Pronase-induced hatching was assessed by incubating embryos at a final pronase concentration of 1 mg/ml at 28 hpf. Hatching time was recorded when all the embryos became free of the chorion. For a separate set of embryos, a normal hatching event was also recorded. The hatching event was considered complete once the embryos were free of the chorion.

### 2.3 Whole-mount in situ hybridisation (WISH) and immunohistochemistry (IHC)

#### 2.3.1 WISH Probe synthesis

Sequences for hatching gland markers *cathepsin Lb* (*ctslb*) and *hatching enzyme 1, tandem duplicate 1* (*he1.1*) were amplified using the following primers:

A. *ctslb*-forward primer: AGACCGCCTCTATGTTCGGA; reverse primer: **TAATACGACTCACTATAGGG**AGCGACATTAAAACGGGGGT (The bold part in the reverse primer is the T7 promoter sequence).
B. *he1.1*-forward primer: GGAGATGTGGTGCTTCCCAA; reverse primer: **TAATACGACTCACTATAGGG**ACAGAAATTCCACACAATCACAGA

The PCR products were purified (Clean Up Concentrator Kit, #021-250C, A&A Biotechnology) and used as templates to synthesize digoxygenin-labeled antisense RNA probes by in vitro transcription (Dig RNA labelling Mix, #11277073910, Roche; MAXIscript T7 transcription kit, #AM1314, Invitrogen). WISH was performed at various developmental stages using an established protocol (40) with slight modifications.

#### 2.3.2 IHC

Embryos were fixed in 4% PFA/PBS for 2 hours at room temperature, followed by washing with PBS (2×1 min; then 3×10 min). For antibody penetration, embryos were briefly treated: 10 seconds in MilliQ water, 8-10 min in ice-cold acetone, and again 10 seconds in MilliQ water. Immediately following this treatment, embryos were thoroughly washed with PBS (3×15 min) at room temperature. Embryos were then blocked for 2 hours at room temperature using 10% Bovine serum albumin (Sigma) diluted in PBST (PBS containing 0.1% Tween 20).

Primary antibodies: rabbit polyclonal anti-Ctsl1b (#GTX128324, GeneTex) and rabbit polyclonal anti-phospho histone H3 (Ser-10) mitosis marker (#06-570, Sigma) were diluted 1:200 in blocking solution overnight at 4°C. Donkey anti-rabbit Alexa Fluor 488 (1:1000, #R37114, Thermo Fisher, USA) was used as a secondary antibody. Special care was taken throughout both procedures due to the extreme sensitivity of HG cells. Images were captured using a Nikon stereo microscope (SMZ25). To quantify the number of HG cells, the Cell Counter plugin in ImageJ (Fiji) was used.

### 2.4 Quantitative real-time polymerase chain reaction

To quantify the expression of hatching gland markers, qRT-PCR was performed at 28, 50, and 76 hpf. TRIzol (#T9424, Sigma) -chloroform (#2344331116, POCH) method was used for isolating RNA (50 whole embryos) following the manufacturer’s protocol. One microgram of quantified RNA was used to synthesize cDNA (iScript cDNA synthesis kit, #1708890, Bio-Rad). qRTPCR was performed as described (34). A complete list of genes with primer sequences is provided in Table 1.

**Table 1:**
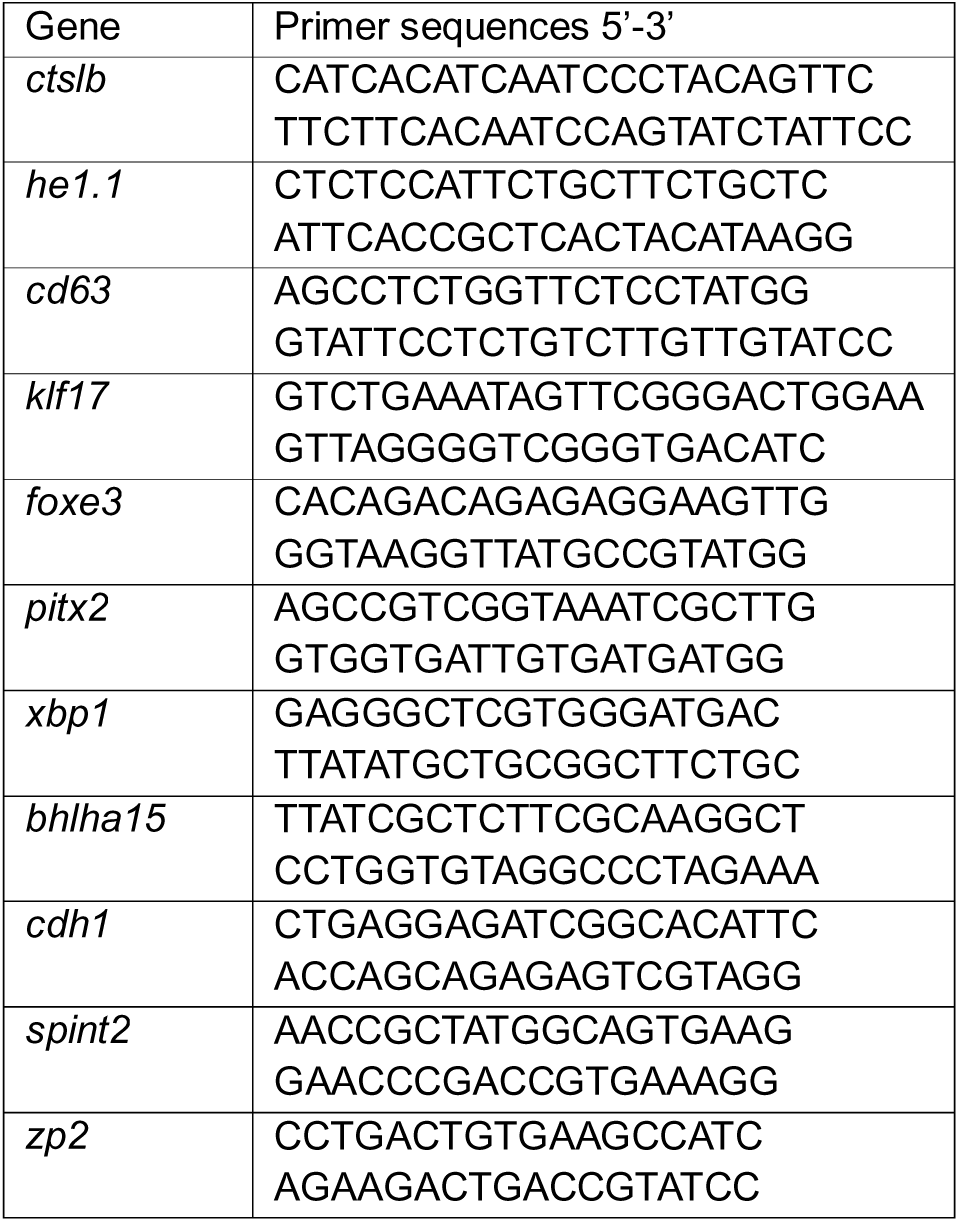
List of qRT-PCR primers

### 2.5 Western blot

Protein lysates were prepared using 50 embryos at 28, 50, and 76 hpf. Thirty micrograms of lysate per lane was separated on a 10% polyacrylamide gel (90 volts) and transferred to a PVDF membrane (#1620177, Bio-Rad). The membrane was blocked with 5% milk and incubated overnight with primary antibodies: rabbit polyclonal anti-Ctsl1b (1:5000, #GTX128324, GeneTex) and mouse monoclonal anti-GAPDH (1:1000 in 1% milk, #ZG003, Invitrogen) as an internal control. Goat polyclonal anti-rabbit IgG (#A9169, Sigma) and rabbit polyclonal anti-mouse IgG (A9044, Sigma) were used as secondary antibodies at 1:5000 dilution each. Protein detection was performed using chemiluminescence and visualized with a Gel Doc system (Bio-Rad).

### 2.6 Enzyme secretion assay

Proteolytic activity (protease secretion) was quantified using the Protease Fluorescent Detection Kit (#PF0100, Sigma). At 34-36 hpf, 30 manually dechorionated embryos per group were individually placed in a 96-well plate containing 100 μL of E3 medium and incubated at 28.5°C. At 50 hpf, 90 μl of medium from 20 wells/group was pooled and concentrated using a 10 kDa ultra centrifuge unit (#UFC5010, Amicon) (7). Later, proteolytic activity was quantified according to the manufacturer’s protocol (fresh E3 medium served as a negative control; water as blank; Trypsin as positive control). Fluorescent intensity was measured using a Tecan instrument at an excitation wavelength of 485 nm and emission wavelength of 535 nm.

### 2.7 FM 1-43 staining for visualizing exocytosis

*In vivo* visualization of exocytosis was performed at 65 hpf using the lipophilic styryl dye FM 1-43 (*N*-(3-Triethylammoniumpropyl)-4-(4-(Dibutylamino) Styryl) Pyridinium Dibromide). PTU-treated, manually dechorionated embryos were anesthetized (MS222) and incubated for 30 seconds in 3 μM FM 1-43 (#T3163, Invitrogen) in E3 medium. Following three rinses in fresh E3 medium, embryos were embedded in 1.5% LMA in E3 medium (supplemented with 0.02% tricaine) and imaged using the 20x objective of a Zeiss LSM800 confocal microscope with the dedicated FM 1-43 laser. Maximum-intensity projections of each z-stack were generated, and the images were processed using the ZEN software.

### 2.8 Live imaging

To visualize the morphology and granularity of HGCs, live imaging was performed. At 76 hpf, PTU-treated embryos were manually dechorionated, anesthetized using MS222/tricaine (0.02%, Sigma), and mounted on a capillary using 1.5% low-melting agarose (LMA, #50080, Lonza; prepared in E3 medium containing 0.02 % tricaine). High-resolution images of the zebrafish HG cells were acquired using transmitted LED illumination on a Zeiss Lightsheet Z1 microscope equipped with a 40x objective. Images were subsequently processed using the ZEN (Zeiss) software.

### 2.9 Cell death analysis via TUNEL assay

To assess apoptosis-induced cell death in HG cells, a TUNEL (terminal deoxynucleotidyl transferase-mediated dUTP nick end labeling) assay was performed at 18 hpf using an *in situ* cell death kit (Fluorescein, #11684795910, Roche) according to the manufacturer’s protocol. Images were obtained and processed similarly to the previous section using a confocal microscope (LSM800).

### 2.10 Chemical treatments with potassium and 4-aminopyridine

The effect of a potassium channel blocker on hatching was evaluated. At the 2-8 cell stage, 12 embryos per group were incubated in varying final concentrations of 4-aminopyridine (4-AP, 1-4 mM/L, #A78403, Sigma) diluted in E3 medium. Embryos were maintained at 28.5°C, and the hatching event was recorded as mentioned earlier.

### 2.11 Statistical analysis

Statistical analysis for hatching assessment, HG cell count, and proteolytic activity was performed using GraphPad Prism (Version 9.5.1). The Shapiro-Wilk test was used to determine the normality of the data. Comparisons were made using one-way ANOVA followed by a post hoc test. A p-value of ≤0.05 was considered statistically significant.

For gene expression analysis, fold change in the mutants relative to wild-type was calculated using the delta-delta-C(t) method, with eef1a1l1 (eukaryotic translation elongation factor 1 alpha 1-like *1*) as an internal control. Statistical significance was determined using one-way ANOVA.

In all analyses, a p-value of ≤0.05 was considered statistically significant.

## 3 RESULTS

### 3.1 Kv2.1 channel subunit mutations affect hatching dynamics

We first examined the effect of the *kcnb1* and *kcng4b* mutations on the hatching process.

#### 3.1.1 Pronase-induced hatching

Hatching time was assessed after pronase treatment at 28 hpf. Interestingly, *kcnb1* mutants exhibited accelerated hatching, completing the process within 12 minutes of incubation, compared to WT embryos (∼15 min) (Fig. 1A). In contrast, *kcng4b* mutants showed delayed hatching, requiring 20-22 minutes.

**Fig. 1:**
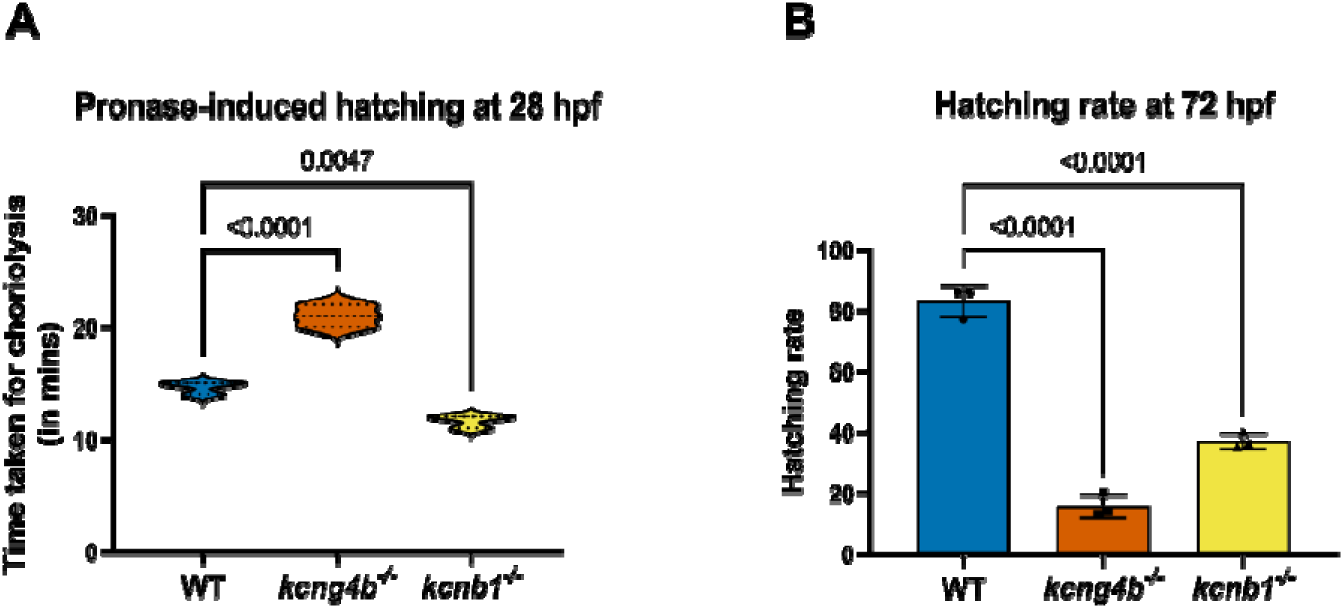
Mutations in the Kv channel subunit alter zebrafish embryonic hatching time. A) Pronase-induced hatching assay: Embryos were treated with pronase (1 mg/ml at 28 hpf) to induce chorion softening exhibited differential responses: *kcng4b* mutants showed a significant delay in hatching time, whereas *kcnb1* mutants displayed accelerated hatching compared to the WT controls. Data are represented as a violin plot. N=3; n = n WT=450, n *kcng4b^-/-^=*450*, n kcnb1^-/-^ =*440 (total time taken for around 145-150 embryos/group to hatch was recorded per replicate). Statistics were calculated using one-way ANOVA followed by Dunnett’s multiple comparison test. A p-value of ≤ 0.05 is considered significant. B) Normal hatching rate: The percentage of hatched embryos recorded at 72 hpf was significantly reduced in both *kcnb1* and *kcng4b* mutants compared to WT. Data are represented as Mean ± SD. N=3; n WT=110, n *kcng4b-C1^-/-^=*100*, n kcnb1^-/-^=*80 (Overall hatching rate was recorded for 25-40 larvae per group per replicate). Statistics were calculated using one-way ANOVA followed by Dunnett’s multiple comparison test. A p-value of ≤ 0.05 is considered significant.

#### 3.1.2 Normal hatching

To assess the overall effect of mutation on hatching, the embryos were tracked until hatching was complete. While 80% of WT embryos had hatched by 72 hpf, this proportion was less than half in both mutants. The *kcng4b* mutants showed severe delay, with only 20% hatched by this time point (Fig. 1B). All mutant embryos eventually completed hatching by 96 hpf. This altered hatching phenotype suggests a role for Kv2.1 channel subunits in regulating zebrafish hatching.

### 3.2 Abnormal HG patterning and reduction of HG cells

Next, we performed WISH for the HG markers cathepsin Lb (*ctslb*) and hatching enzyme 1, tandem duplicate 1 (*he1.1*) to visualize the HG morphology and development. Early expressions in the polster (9-11 hpf) showed only subtle differences in the expression pattern between the wild-type and the mutants (Supplementary Fig.1). However, as the HG cells migrated, severe patterning defects emerged in both *kcng4b* and *kcnb1* mutants. At 14 hpf, after migration from the polster, an abnormal patterning of the HG characterized by reduced expression of *ctslb* and *he1.1* was observed in the mutants (Fig. 2A). The typical necklace-like pattern of hatching gland cells on the enveloping layer at 18 hpf was affected, showing gaps in the HG and its narrowing (Fig. 2B). Around 28-30 hpf, an abnormal patterning persisted, marked by the gaps without marker expression. Both mutants also showed a reduction of HG cell layers (Fig. 2C). By 50 hpf, the HG appeared narrower in both mutants (Fig. 2D).

**Fig. 2:**
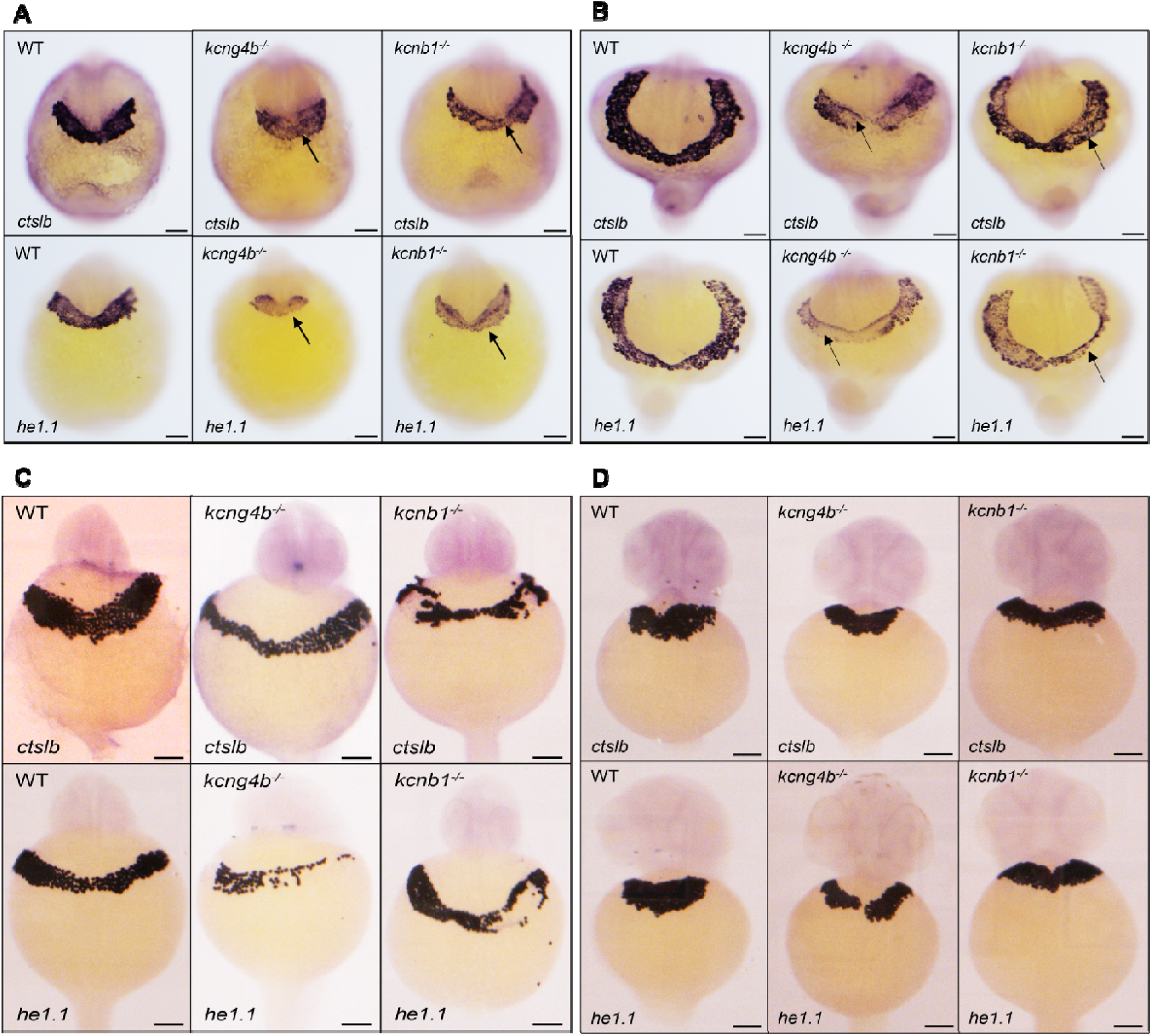
Kv2.1 subunit mutants exhibit progressive defects in the Hatching gland (HG) patterning and morphology. Whole-mount *in situ* hybridization (WISH) was performed using *ctslb* (*cathepsin LB*, top row) and *he1.1* (*hatching enzyme 1, tandem duplicate 1*, bottom row) probes. To visualize HG development from 14-50 hpf. N=3; total number of embryos for all stages WT=15, n *kcng4b^-/-^=*12*, n kcnb1^-/-^=*12. Scale bar is 100 µm for all images. A) 14 hpf (early patterning): While WT shows a continuous region of expression, the expression pattern of *ctslb* and *he1.1* in both mutants was notably affected (indicated by black arrows), suggesting an early developmental defect. B) 18 hpf (migration): WT embryo shows HG cells organized in a cohesive, necklace-like pattern along the anterior enveloping layer. In both mutants, this organization is severely disrupted (black arrows), indicating failure in coordinated cell migration and/or adhesion. C) 28 hpf (Morphogenesis): In WT embryos, HG cells have consolidated to form a thick, uniform band. In contrast, both *kcng4b* and *kcnb1* mutants exhibit narrowing of the HG, evidenced by reduced overall tissue width. D) 50 hpf (Pre-hatching stage): WT HG cells show a mature morphology, clustering, and migrated anteriorly, forming a thick tissue mass. The HG tissue in both *kcng4b* and *kcnb1* mutants remains narrower than WT, confirming sustained morphological defect.

The IHC staining using an anti-Ctsl1b antibody confirmed similar patterning defects at 28 hpf (Fig. 3A) and HG narrowing at 50 hpf (Supplementary Fig. 2). Quantification of GFP-positive HG cells (after antibody staining) revealed a significant decrease in the HG cell count in both *kcng4b* and *kcnb1* mutants at 28 hpf (Fig. 3B). These findings collectively indicate that mutations affect the development and patterning of HG.

**Fig. 3:**
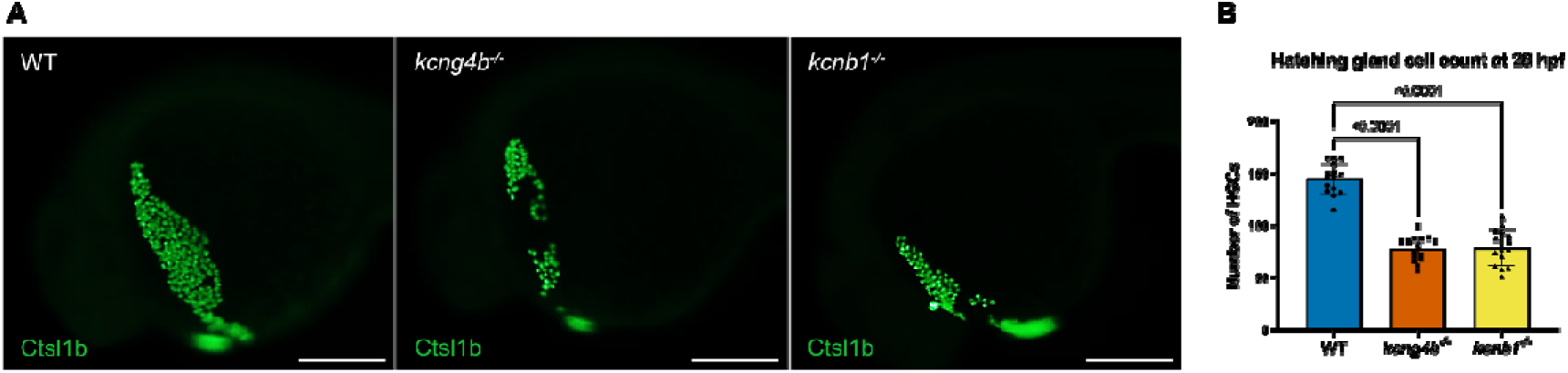
Kv channel mutants show abnormal HG patterning and reduced cell count. A) Cathepsin expression and HG patterning: IHC staining was performed at 28 hpf using anti-Ctsl1b (Cathepsin L1 B) antibody. Compared to WT embryos, both *kcng4b* and *kcnb1* mutants display altered and abnormal patterning of the HG. N=3; total number of embryos 18/genotype. Scale bar is 200 µm. B) Quantification of HG cells: A significant reduction in the number of hatching gland cells was observed in both mutants at 28 hpf (quantified after anti-Ctsl1b staining). Data are represented as Mean ± SD. N=3; total number of embryos counted=15/genotype. Statistics were calculated using one-way ANOVA followed by Dunnett’s multiple comparison test. A p-value of ≤ 0.05 is considered significant.

### 3.3 Altered transcript levels of HG markers and transcriptional factors

To assess the expression of HG’s molecular markers qRT-PCR was performed. Transcript levels of *ctslb* were unaltered at 28 and 50 hpf in both mutants. However, *kcnb1* mutants showed a significant upregulation of *ctslb* at 76 hpf, a change not observed in the *kcng4b* mutant (Fig. 4A). Downregulation of *he1.1* expression was observed in the *kcnb1* mutant at 28 and 50 hpf, but not in *kcng4b*. Conversely, both mutants showed an upregulation in *he1.1* transcript levels at 76 hpf (Fig. 4B).

**Fig. 4:**
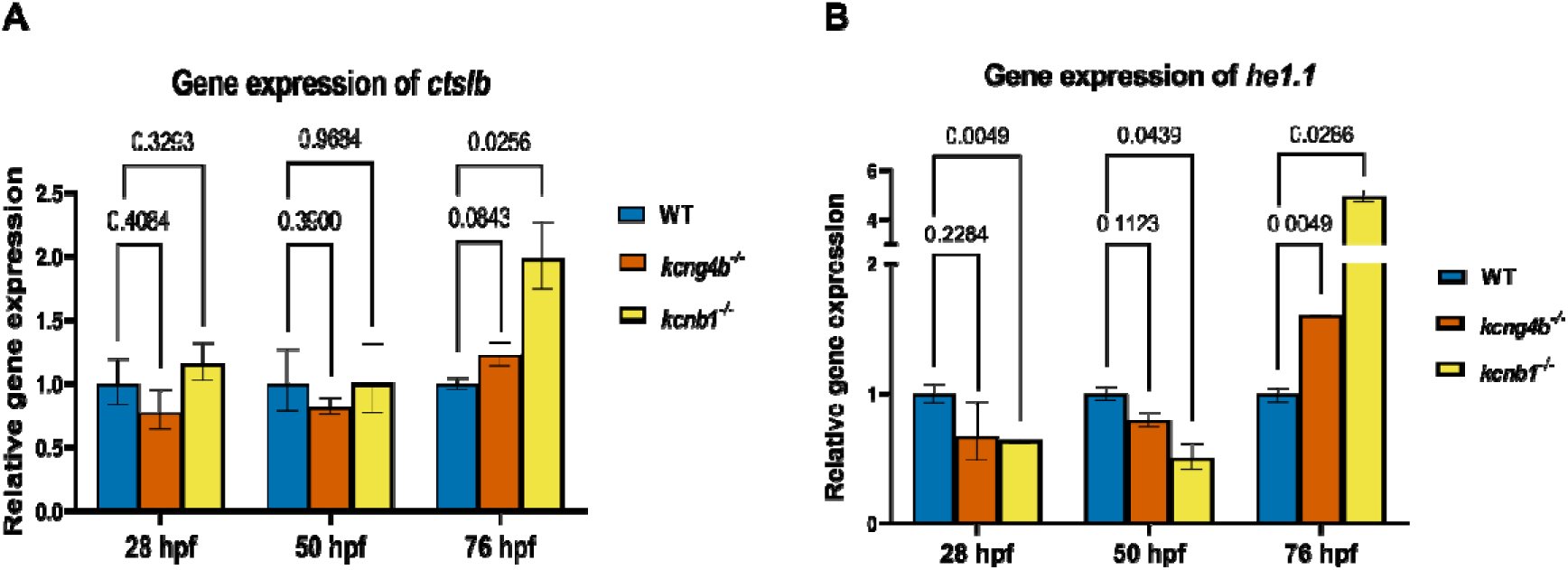
HG marker expression is altered in the Kv channel mutants. qRT-PCR was performed at three key developmental stages (28, 50, and 76 hpf) to analyse the gene expression of HG markers: (A) *cathepsin Lb* (*ctslb*), and (B) *hatching enzyme 1, tandem duplicate 1* (*he1.1).* Fold change in mutants relative to control was calculated using the ΔΔC(t) method. Data are represented as Mean ± SD. N=3; n= 50 pooled larvae/group/replicate. Statistics were calculated by one-way ANOVA. A p-value of ≤ 0.05 is considered significant. Gene expression of *ctslb* was unaltered in both mutants at 28 and 50 hpf. The *kcng4b* mutant showed no difference in the *ctslb* expression at 76 hpf. However, a significant upregulation of *ctslb* expression was observed in the *kcnb1* mutant at 76 hpf. A) *ctslb* gene expression: At 28 and 50 hpf, expression was unaltered in *kcng4b and kcnb1* mutants. Kcnb1 mutants showed a significant upregulation in ctslb expression at 76 hpf, while the expression remained unchanged in the *kcng4b* mutant. B) *he1.1* gene expression: At 28 and 50 hpf, expression was unaltered in both *kcng4b* mutants. Conversely, *kcnb1* mutants showed a significant downregulation of *he1.1*. Both mutants showed a significant increase in *he1.1* transcript level compared to WT.

Moreover, the *kcnb1* mutant showed significant downregulation in the expression levels of several transcriptional factors previously implicated in HG development. *kruppel-like factor 17* (*klf17*), *forkhead box E3* (*foxe3*), and *paired-like homeodomain 2* (*pitx2*) were downregulated at 28 hpf, while in the 28 hpf *kcng4b* mutant, only *pitx2* was mildly downregulated (Supplementary Table 1). Additionally, the expression level of *E-cadherin* (*cdh1*), a gene essential for HG cell migration (12), was significantly downregulated only in the *kcng4b* mutant at 28 hpf. Crucially, the expression levels of secretion-essential transcriptional factors of HG: *X-box binding protein 1* (*xbp1*), *basic helix-loop-helix family, member a15* (*bhlha15*) (41), and HG cell migration/survival gene *serine peptidase inhibitor, Kunitz type, 2* (*spint2*) (12) remained unaltered in both mutants. The expression of *zp2* (chorion marker) was significantly upregulated in both mutants at 28 hpf.

### 3.4 Reduced cathepsin protein synthesis and proteolytic activity

#### 3.4.1 Decreased Cathepsin protein expression

Western blot analysis was performed on whole embryo lysates. While Cathepsin protein levels were similar across all genotypes at 28 hpf (Supplementary Fig. 3), an apparent decrease in cathepsin levels was observed in both mutants at 50 hpf. At 76 hpf, cathepsin protein levels were very low, resulting in faint bands in all samples (Fig. 5A). This suggested reduced cathepsin synthesis in the mutants.

**Fig. 5:**
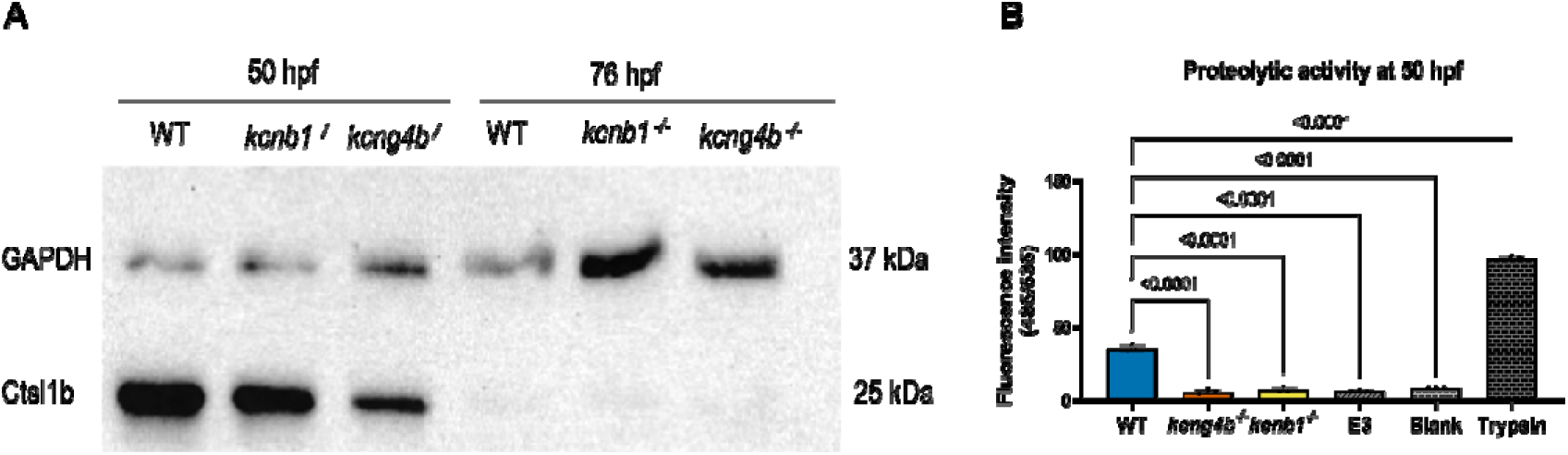
Reduced cathepsin protein synthesis and proteolytic activity. A) Western Blot Analysis: Whole embryo lysates were analysed at 50 and 76 hpf using anti-Ctsl1b (Cathepsin L1 B) antibody to quantify cathepsin synthesis. Cathepsin protein levels were reduced in the *kcnb1* and *kcng4b* mutants at 50 hpf. By 76 hpf, cathepsin expression was nearly absent in all three (WT, *kcng4b^-/-^*, and *kcnb1^-/-^*). GAPDH was used as a loading control. N=3; n= 50 pooled larvae/group/replicate. B) Quantification of proteolytic activity: Proteolytic activity of hatching enzymes was quantified from the medium at 50 hpf using a protease fluorescent detection kit (casein substrate). Both mutants showed significantly reduced proteolytic activity, comparable to the E3 and blank controls. Trypsin was used as a positive control. N=3; n= medium from 20 embryos was pooled per group/replicate. Statistics were calculated using one-way ANOVA followed by Dunnett’s multiple comparison test. A p-value of ≤ 0.05 is considered significant.

#### 3.4.2 Reduced hatching enzyme activity

We assessed secretion of hatching enzymes by measuring proteolytic activity in E3 medium collected from dechorionated embryos at 50 hpf. The fluorescence intensity, which is directly proportional to the protease activity, was significantly lower in the media from both *kcng4b* and *kcnb1* mutants compared to the wild-type. Activity levels in the mutant media were comparable to the E3 medium and the blank (Fig. 5B). This result confirms that proteolytic activity is markedly reduced in the Kv channel mutants.

### 3.5 Mutation in Kv channel subunits results in delayed secretion of hatching enzymes

#### 3.5.1 Exocytosis visualization

To visualize the underlying cellular process, we performed *in vivo* exocytosis staining using the FM 1-43 styryl dye at 65 hpf. This dye is internalized upon vesicle recycling post-exocytosis (42). Wild-type embryos showed active exocytosis, evident by numerous bright orange recycled vesicles inside the HG cells (Fig. 6A). In sharp contrast, Kv2.1 mutants displayed very few bright orange dots, suggesting reduced secretion of hatching enzymes.

**Fig. 6:**
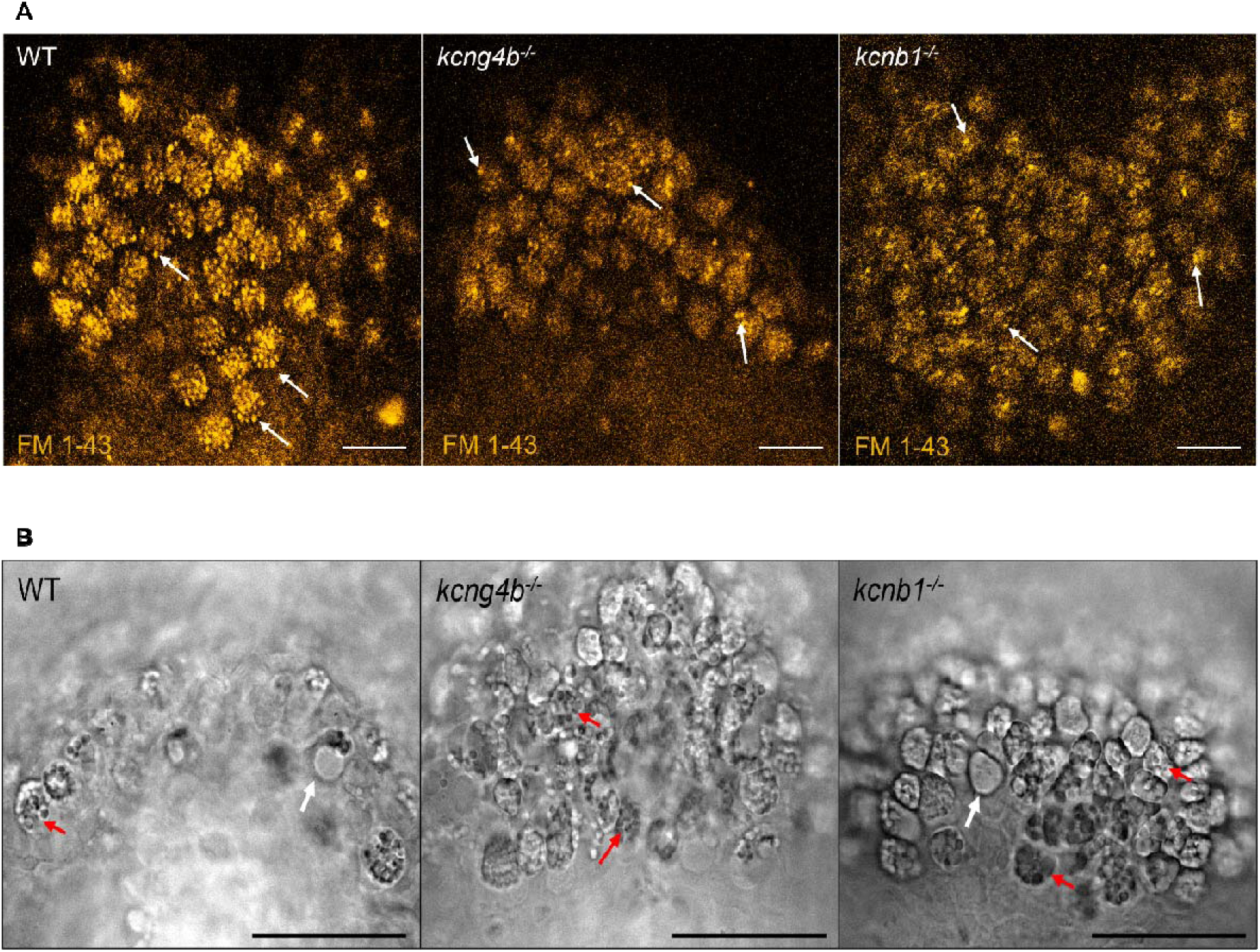
Kv2.1 subunit mutants exhibit defects in terminal hatching enzyme secretion. A) Visualization of active exocytosis using FM 1-43 dye (65 hpf): The lipophilic styryl dye FM 1-43 (*N*-(3-Triethylammoniumpropyl)-4-(4-(Dibutylamino) Styryl) Pyridinium Dibromide) was used for visualizing active exocytosis, as the dye internalizes upon vesicle recycling post secretion. WT embryos exhibited high active exocytosis in the HG, characterized by numerous bright orange dots (indicated by white arrows), reflecting robust enzyme secretion. In contrast, *kcng4b* and *kcnb1* mutants display significantly reduced exocytosis, with only a few FM 1-43 positive dots observed. N=3; n=9/genotype. Scale bar is 20 μm. B) Live Imaging of HG cells (76 hpf): The zebrafish HG undergoes holocrine secretion, where the cell ruptures and dies upon enzyme release. At 76 hpf, WT embryos show only a few HG cells packed with secretory granules (indicated using red arrows). This is consistent with successful holocrine secretion and hatching. Conversely, both *kcng4b* and *kcnb1* mutants retain numerous HG cells still densely packed with granules, demonstrating a severe defect in the terminal secretion. N=3; n=9/genotype. Scale bar is 50 μm.

#### 3.5.2 HG cell density at 76 hpf

Live imaging of zebrafish embryos at 76 hpf revealed that wild-type embryos had fewer secretory granule-packed HG cells (Fig. 6B). An empty cell with no secretory granule is also visible, and this aligns with the HG cell’s holocrine secretion mechanism, where the cell dies upon enzyme release. The mutants, however, retained a higher number of HG cells that were densely packed with secretory granules, indicating a delay in the release of hatching enzymes.

Together, these findings strongly suggest that the mechanism underlying hatching delay is the delayed secretion of hatching enzymes in the mutants of Kv channel subunits.

#### 3.6.1 Apoptosis (TUNEL)

To determine if the observed reduction in the HG cell number was due to apoptosis, we performed a TUNEL assay at 18 hpf. While numerous apoptotic cells are observed throughout the embryos, they are more concentrated near the tail. However, a few apoptotic cells were observed in the HG of both *kcng4b* and *kcnb1* mutants (Supplementary Fig. 4), suggesting that mutations in Kv2.1 subunits induce early apoptosis.

#### 3.6.2 Cell proliferation (pHH3)

IHC staining for cell proliferation marker anti-phospho histone H3 (pHH3) showed no proliferative cells within the gaps in the mutant HG (Fig. 7).

**Fig. 7:**
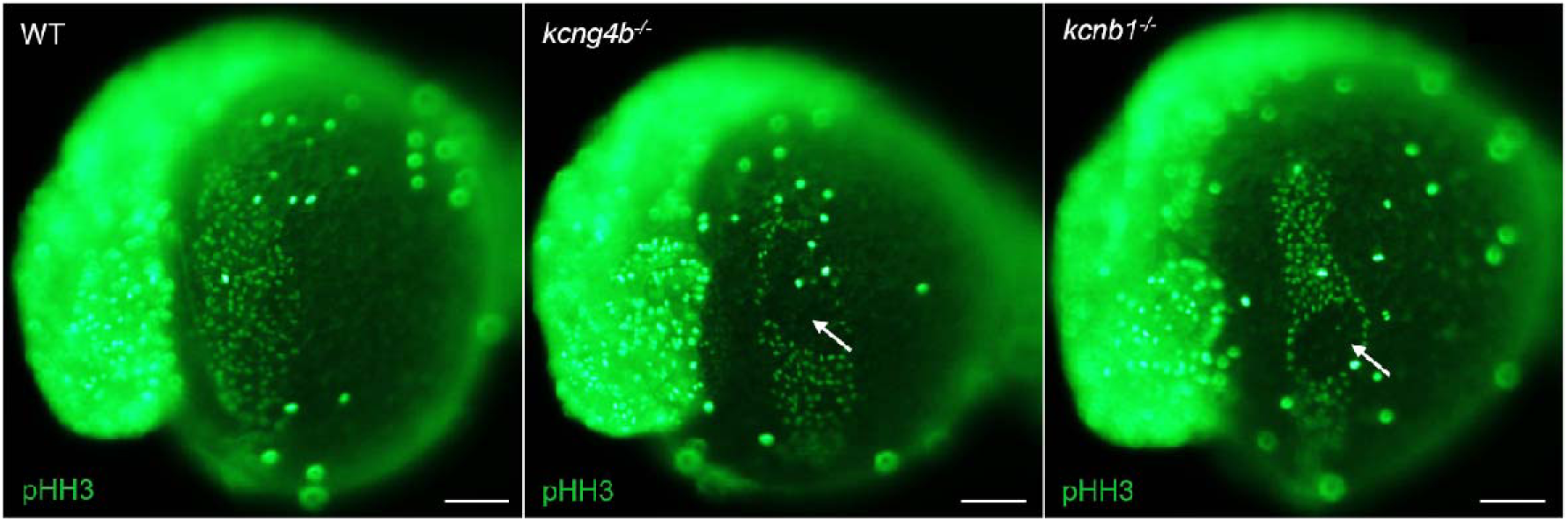
Cell proliferation defects in the HG of Kv2.1 subunit mutants. Visualization of proliferative nuclei using anti-pHH3 (24 hpf): IHC staining was performed using anti-pHH3 (phospho-histone H3) antibody to visualize mitotic cells. Compared with WT embryos, the HG of both *kcng4b* and *kcnb1* mutants showed compromised patterning, evidenced by the absence of proliferative cells within the gapped regions (indicated by white arrow). N=3; n=12/genotype. Scale bar is 100 μm.

#### 3.7.1 Survival and toxicity

We first assessed the general toxicity of 4-AP in all three genotypes. The lowest concentration (1 mM) was tolerated by WT and *kcnb1* mutant embryos throughout the experiment, although the survival of *kcng4b* mutant embryos was significantly compromised at day 3 (Supplementary Fig. 5A). At 2 mM, *kcng4b* mutants died at day 2, whereas WT and *kcnb1* mutant embryos survived (Supplementary Fig. 5B). Treatment with a higher concentration (4 mM) of 4-AP, proved lethal to *kcng4b* mutant embryos by day 1, whereas WT and *kcnb1* mutant embryos showed about 50-60% survival, but by day 2, they also died (Supplementary Fig. 5C). These results indicated a hypersensitivity of the *kcng4b* mutant to the global inhibition of Kv channel.

#### 3.7.2 Hatching rescue

The delayed hatching phenotype could be due to a defect in membrane potential. Thus, we tested whether pharmacologically induced plasma membrane depolarization could affect HG development in mutants. Treatment with 1 mM 4-AP rescued the delayed hatching phenotype in both *kcnb1* and *kcng4b* mutants, with all embryos from the 1 mM-treated group hatching by the end of day 2 (Fig. 8A). However, hatching was completed in embryos treated with 2 mM of 4-AP on day 3 (Fig.8B). These results confirm that the hatching defect could be attributed to a functional deficiency in the Kv2.1-mediated electrical regulation of the HG cells.

**Fig. 8:**
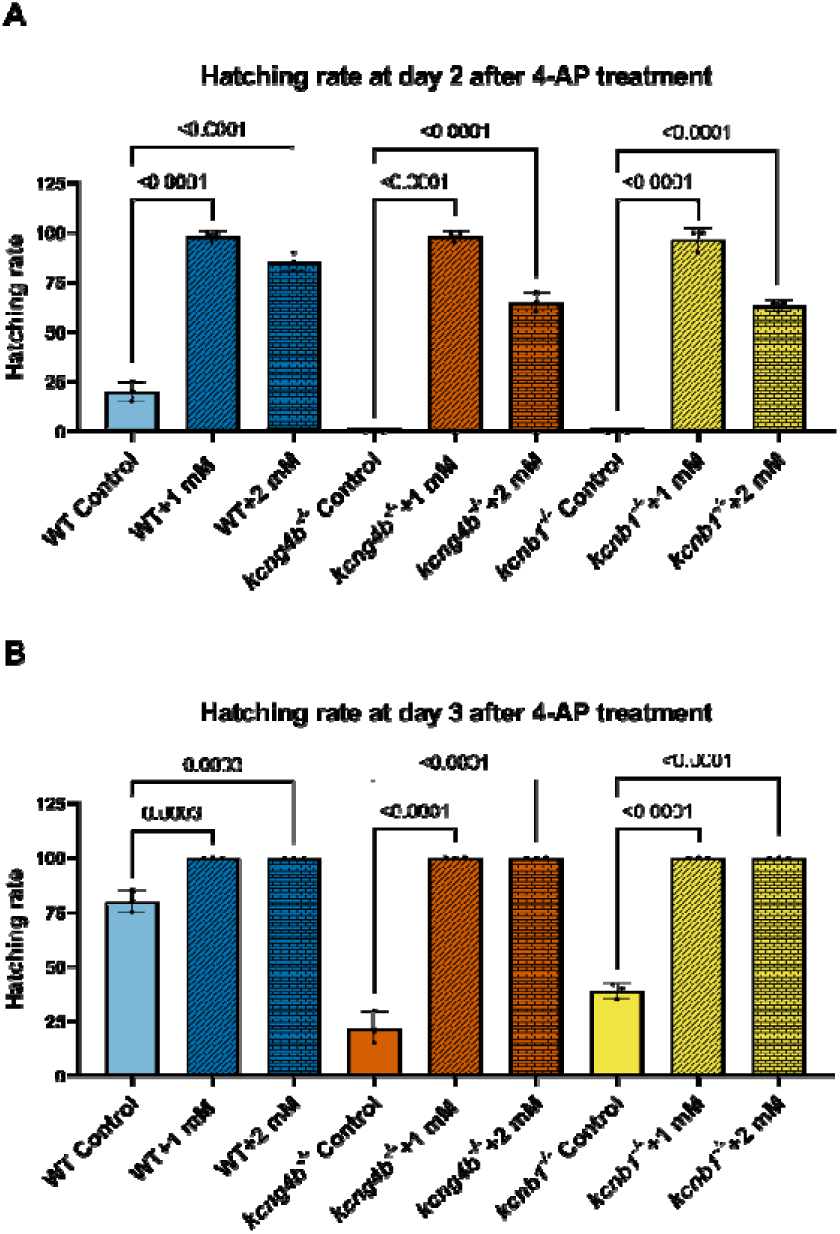
Pharmacological depolarization by 4-aminopyridine (4-AP) rescues the delayed hatching phenotype in the Kv2.1 subunit mutants. Kv channel blocker, 4-aminopyridine (4-AP) was used to depolarize the embryos pharmacologically. Data are represented as Mean ± SD. N=3; total number of embryos 36/genotype/treatment group. Statistics were calculated using one-way ANOVA followed by Dunnett’s multiple comparison test. Control refers to no 4-AP treatment. A p-value of ≤ 0.05 is considered significant. A) Hatching rate at Day 2 (50 hpf): Treatment with 1 mM 4-AP significantly accelerated hatching across all genotypes. By day 2, WT control embryos showed a moderate hatching rate (∼23%), but both kcng4b and kcnb1 control embryos exhibited a near-zero hatching rate. Treatment with 1 mM 4-AP completely overrides the Kv2.1 deficiency, resulting in an accelerated hatching rate (∼97%) in all genotypes. 2 mM 4-AP treatment also accelerated hatching, but at different rates (80% in WT; 65% in mutants). B) Hatching rate at Day 3 (75 hpf): Untreated WT embryos completed hatching by day 3 (∼80%), while both kcng4b and kcnb1 controls show a significant and severe delay (only 22% and 35% hatched, respectively). This delayed hatching is fully corrected (100% hatching) in all genotypes treated with 1 mM or 2 mM 4-AP.

## 4 DISCUSSIONS

In this study, we demonstrated that mutations in the electrically active α subunit Kcnb1 and the modulatory subunit Kcng4b of Kv2.1 impair zebrafish hatching. We found that while *kcnb1* and *kcng4b* mutants both exhibit severely delayed normal hatching and reduced proteolytic activity, they antagonize each other’s activity in the pronase-induced hatching assay. The functional defect is traced to two key flaws: early HG patterning abnormalities and a delay in enzyme secretion, as evidenced by reduced FM 1-43 uptake and retention of secretory granules in HG cells at 76 hpf. The successful chemical rescue of the delayed-hatching phenotype using the Kv channel blocker 4-AP confirmed that the primary defect is of membrane excitability, which seems to regulate this process.

The role of Kv2.1 subunits (Kcnb1 and Kcng4b) in regulating HG development is consistent with their known functions in other embryonic midline-derived tissues. Previous studies on these mutants have established their regulatory roles in the development of other midline-adjacent structures, specifically the zebrafish brain ventricles and Reissner fiber. However, these studies, along with other published works, highlighted a striking functional antagonism between the subunits: the *kcnb1* mutation led to reduced brain ventricles and smaller ears, but increased Scospondin secretion and hypertrophy of Reissner fiber, the *kcng4b* mutation resulted in enlarged ventricles and ears but reduced Scospondin secretion and Reissner fiber hypotrophy (32–35). In contrast, in the current study, we observed functional convergence, where both mutants shared the same key defect - delayed normal hatching, with the only observed antagonism being the differential response to pronase-induced hatching. These shared functional roles align with recent findings on developmental epilepsy, which show that Kcnb1 and Kcng4b exhibit functional convergence in the context of brain hyperexcitability (43).

However, the one instance of antagonism is intriguing because of the mutants’ different responses to pronase-induced hatching. Although the function of pronase is chorion softening without the involvement of hatching enzymes (44), the subtle structural variations in the chorion itself, such as differences in thickness or composition (45), may be responsible for the antagonistic results (accelerated vs. delayed pronase hatching). This could alter the chorion’s permeability or susceptibility to exogenous protease. Since the scope of this study focused on HG cells, further research is required to test the hypothesis that Kv2.1 subunits may regulate chorion integrity.

The observed patterning defects in the HG were both timely and significant. They began around 10 hpf and became evident by 14 hpf, aligned perfectly with the onset of *kcnb1* and *kcng4b* gene expression (32,33,36). This early defect, marked by the narrowing of the HG and creation of large gaps in it, resulted in a significant reduction in the number of HG cells at 28 hpf and likely explains the hatching delay. This suggests a critical role for these subunits in regulating cell proliferation, migration, adhesion, and survival during HG development. The early presence of apoptotic cells in the HG of both mutants, followed by cell proliferation defects, strongly supports the hypothesis that HG defects are caused by cell loss. Kcnb1 is known to induce apoptosis in the brain through various mechanisms (46,47), and *kcnb1* zebrafish mutants have previously shown a loss of proliferative cells in the otic vesicle, indicating that Kcnb1 plays a conserved role in cell survival and proliferation across developmental tissues (33). This function is often attributed to Kv2.1’s non-canonical signaling roles, such as its association with integrins to form the Integrin-K^+^ channel (IKC) complex, which regulates migration, proliferation, and cell survival (47). Indeed, mutations in KCNB1 have been shown to impair neuronal migration, causing developmental epilepsy (48). Therefore, the HG patterning defects observed in this study represent one more case in which Kcnb1 and its partner Kcng4b regulate tissue morphogenesis and cell fate, linking membrane excitability to developmental timing.

The specific molecular defects observed strengthen this notion. In the *kcnb1* mutant, the broad downregulation of early HG transcription factors (*klf17, pitx2*, and *foxe3*) suggests a fundamental defect in the specification or differentiation pathway of the HG cells. Conversely, the *kcng4b* mutant showed a decrease in the cell migration factor, *cdh1,* at 28 hpf. This is highly relevant because HG patterning requires complex, coordinated cell migration, and the loss of *cdh1* in *kcng4b* mutants parallels the *spint2* mutant phenotype. *spint2* is described as essential for HG cell survival and migration alongside *cdh1*, and zebrafish *spint2* mutants show complete HG loss by 44-48 hpf (12). Our data did not show changes in *spint2* transcript levels. This divergence indicates that *kcng4b* may act upstream of *cdh1* in regulating migration, resulting in a partial HG defect that is less severe than the total loss observed in *spint2* mutants.

Beyond development, the core finding is the role of these channels in HG functions. Despite not observing any changes in the transcript levels of essential HG-specific secretion factors, *xbp1* or *bhlha15* (41), we observed a significant reduction in overall proteolytic activity in the medium of both mutants at 50 hpf, indicating that less enzyme was secreted. This functional deficiency is immediately supported by the decrease in cathepsin protein synthesis observed at 50 hpf via western blotting, suggesting a late-stage failure in producing or stabilizing the enzyme. Crucially, although we observed an increase in *ctslb* and *he1.1* transcript levels at 76 hpf, this transcriptional upregulation did not translate into increased protein levels. This suggests that HG cells, despite potentially being fewer due to earlier HG cell loss, attempt to compensate transcriptionally, but fail to execute the timely final secretion. Consistent with the HG’s holocrine secretion mechanism (16), the definitive proof of a secretory delay comes from the reduced exocytosis (FM 1-43 staining) and retention of granule-packed HG cells at 76 hpf in both mutants, demonstrating a direct functional failure in the final secretion step. As Kv2.1 is well known to regulate secretion in other specialized tissues, such as insulin secretion (49–51), chromaffin granule release (52), and negative regulation of Scospondin production (35), this novel role in HG secretion is consistent with its established function as a modulator of specialized secretory cell function.

Extending this regulatory role to other specialized, terminally differentiated cells, Kcng4 loss causes male sterility in rodents by disrupting spermiogenesis, a final maturation step sensitive to membrane potential regulation (30). A similar reduction in fertility has been observed in the dominant-negative *kcng4b^waw305^* allele (35,39). In contrast, no apparent changes in fertility were detected in Kcnb1 KO mice, which mimic a human mutation affecting voltage sensing (53). Perhaps the nature of the genetic defect may play a role in shaping the mutant phenotype. The mutations analyzed in this study were shown to affect the expression level of mutated genes (43,54). The delayed hatching phenotype and disruption of terminal holocrine secretion in the zebrafish mutants seem to indicate a conserved need for the Kv2.1 complex in regulating the timing and function of specialized terminally differentiated cells in both fish and mammals. This is particularly relevant when considering the equivalent process of human blastocyst hatching, in which terminally differentiated, specialized trophoblast epithelial cells (zona breakers) secrete proteases required for blastocyst hatching (5). Our data suggest that Kv channel regulation of enzymatic secretion is a key evolutionary mechanism for crucial life-stage transitions.

We propose that Kcnb1 and Kcng4b subunits are required to precisely modulate the membrane potential of HG cells, facilitating the significant and sustained Ca^2+^ influx necessary to trigger both vesicle fusion (exocytosis) and terminal cell rupture for holocrine release of hatching enzymes. This physiological model is strongly supported by the chemical rescue experiment: the delayed hatching phenotype was alleviated by 4-AP, a Kv channel blocker. In typical secretory cells, Kv channel blockers, such as 4-AP, cause depolarization of the membrane, prolonging the open time of voltage-gated Ca^2+^ channels, thereby enhancing calcium influx (55). This calcium influx helps the mutants overcome the intrinsic Kv channel deficit. Thus, Kv2.1 subunits act as crucial controllers of the electrical state of HG cells, regulating the final Ca^2+^-mediated secretory event. While acknowledging that 4-AP may also increase larval movement, potentially contributing to the mechanical component of the rescue, the successful alleviation of the hatching delay strongly implicates a defect in membrane potential regulation. Therefore, the delayed hatching phenotype results from a two-pronged failure: early developmental defects that lead to fewer HG cells, compounded by a late-stage functional defect in the enzyme release mechanism.

Our study identifies Kcnb1 and Kcng4b as functionally essential and parallel components in regulating zebrafish hatching. Their loss results in a cascade of defects, starting with impaired HG development and patterning, leading to a dramatic reduction and delay in hatching enzyme secretion. This study highlights Kv2.1 subunits as necessary timekeepers in embryonic development, required for the precise regulation of HG holocrine secretion. Further studies should focus on the specific Ca^2+^ dynamics within the mutant HG cells to fully understand how these two Kv channel subunits coordinate the final secretory event.

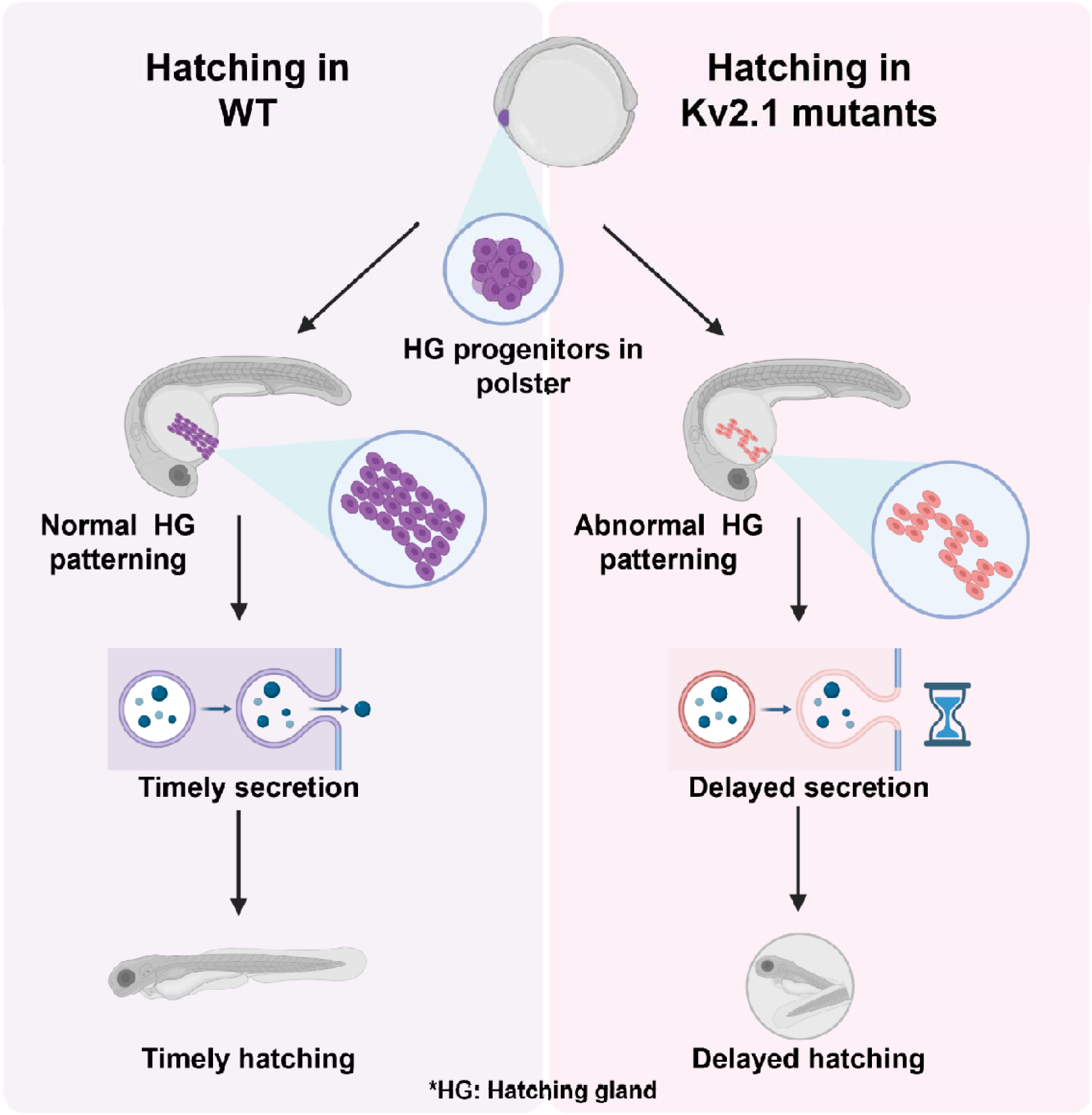

## FUNDING DECLARATION

Korzh V acknowledges support from the Opus grant of the National Science Centre (NCN), Poland (2020/39/B/NZ3/02729).

## CONSENT TO PUBLISH

Not applicable.

## CONSENT TO PARTICIPATE

Not applicable.

## ETHICS DECLARATION

*Danio rerio* lines were maintained following established standard husbandry protocols (38) in the Zebrafish Core Facility of the International Institute of Molecular and Cell Biology, Warsaw (license No. PL14656251). All procedures involving embryos and larvae adhered to the guidelines of the Polish Laboratory Animal Science Association and European Communities Council Directive (63/2010/EEC).

## DATA AVAILABILITY DECLARATION

The materials and reagents described in the paper are available from the authors upon request.

## AUTHOR CONTRIBUTION DECLARATION

Ruchi P. Jain: Conceptualization, Methodology, Investigation, Writing – original draft, Project administration, Data curation. R. Rosa Amini: Methodology, Investigation. Vladimir Korzh: Writing – review & editing, Validation, Supervision, Project administration, Funding acquisition.

## DECLARATION OF GENERATIVE AI AND AI-ASSISTED TECHNOLOGIES IN THE WRITING PROCESS

The authors used Grammarly to enhance the language and readability of the manuscript. Authors confirm that all generated and edited content was rigorously reviewed and validated, and they assume full responsibility for the final version of this publication.

## ACKNOWLEDGEMENTS

We are deeply grateful to Prof. Jacek Kuznicki for his continuous support and invaluable guidance. We also wish to acknowledge the valuable discussions with all members of the Laboratory of Neurodegeneration (IIMCB in Warsaw). Finally, we are grateful to the Microscopy and Zebrafish Core Facilities (IIMCB in Warsaw) for their expert technical assistance and dedicated fish maintenance.

## COMPETING INTEREST DECLARATION

Authors declare no competing interests.

## Notes

### Competing Interest Statement

The authors have declared no competing interest.

