## Supplemental Figure 1 for "From patterning to secretion: Kv2.1 subunits as regulators of zebrafish hatching gland morphogenesis and function"

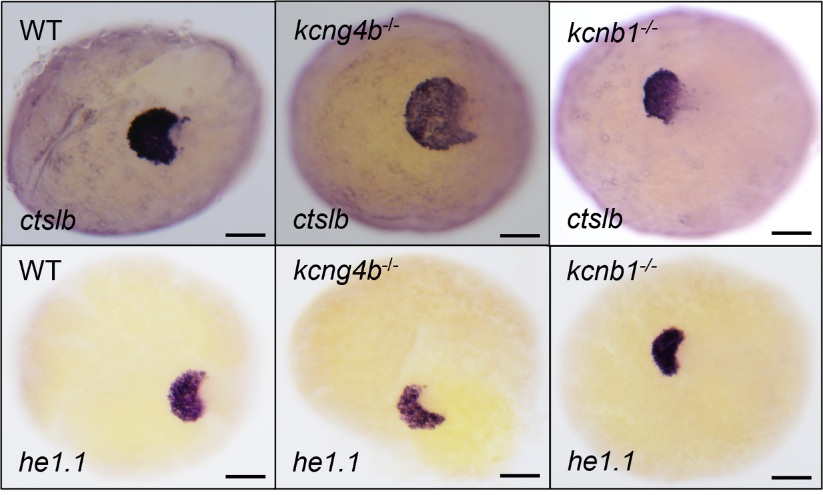
**Supplementary Fig. 1: Early hatching gland (HG) patterning is affected in the Kv channel mutants (9-11 hpf)**

Whole-mount *in situ* hybridization (WISH) was performed using HG markers *ctslb* (*cathepsin LB*, top row) and *he1.1* (*hatching enzyme 1, tandem duplicate 1*, bottom row) to visualize HG development at 9-11 hpf. While WT embryos exhibit consistent expression for both markers, the expression of ctslb is altered (reduced expression) in the kcng4b and kcnb1 mutants. N=3; total number of embryos 12/genotype. Scale bar is 100 µm for all images.

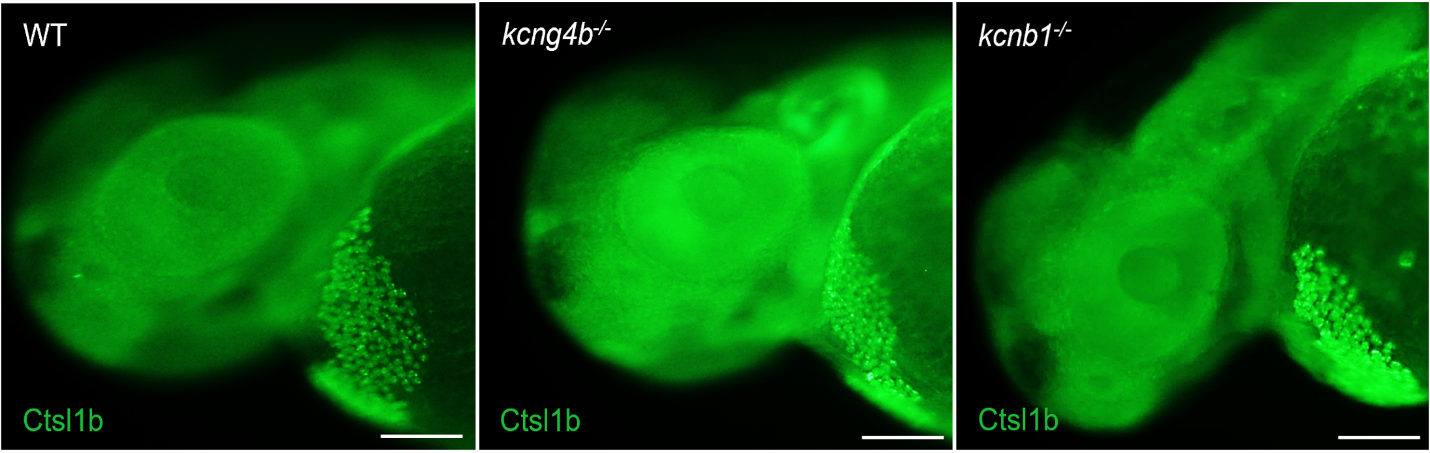
**Supplementary Fig. 2: Kv channel mutants show abnormal HG patterning**

IHC staining was performed at 50 hpf using anti-Ctsl1b (Cathepsin L1 B) antibody. Compared to WT embryos, both *kcng4b* and *kcnb1* mutants display altered and abnormal patterning of the HG, evident by a narrower HG in the mutants compared to the WT. N=3; total number of embryos 18/genotype. Scale bar is 100 µm.

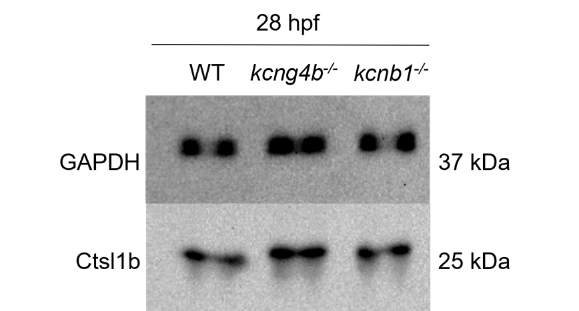

**Supplementary Fig. 3: Cathepsin protein synthesis is unaltered in Kv2.1 subunit mutants at 28 hpf**

Whole embryo lysates were analysed at 28 hpf using anti-Ctsl1b (Cathepsin L1 B) antibody to quantify cathepsin synthesis. Cathepsin protein levels remain unaltered in both *kcng4b* and *kcnb1* mutants. GAPDH was used as a loading control. N=3; n= 50 pooled larvae/group/replicate.

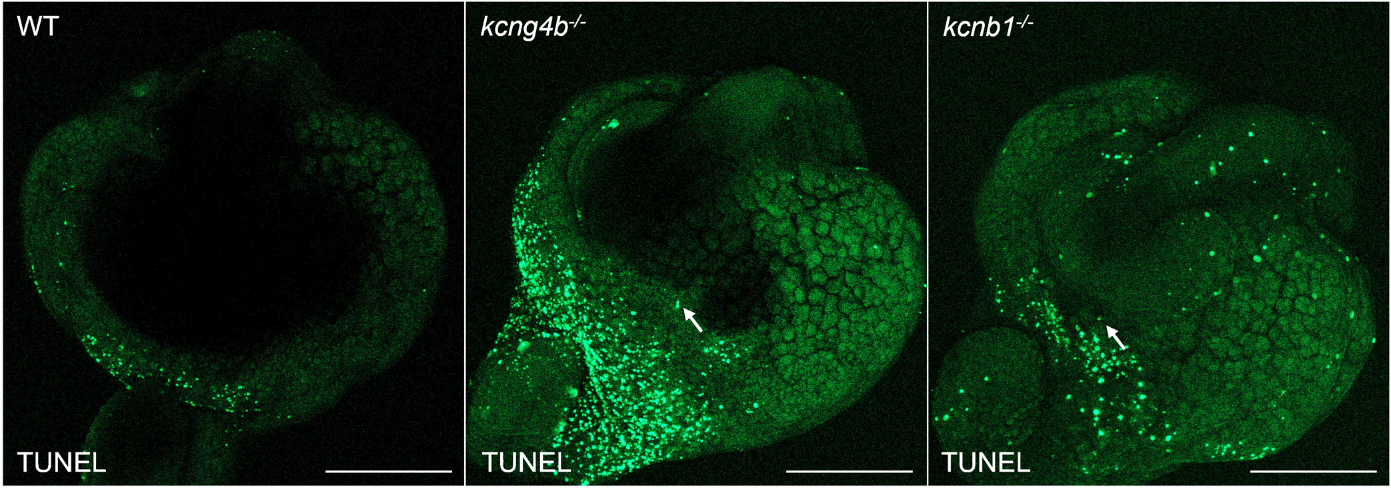

**Supplementary Fig.4: Apoptosis observed in the HG of Kv2.1 subunit mutants**

*In situ* cell death was visualized using the TUNEL (terminal deoxynucleotidyl transferase-mediated dUTP nick end labeling) assay kit. The assay revealed the presence of a few apoptotic cells (indicated by white arrows) within the HG of both *kcng4b* and *kcnb1* mutants. N=3; n=6/genotype. Scale bar is 200 μm.

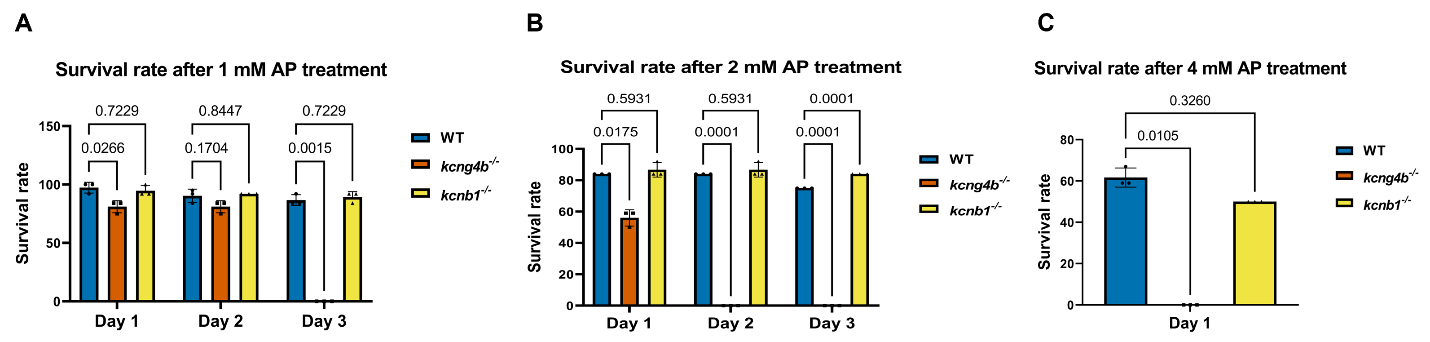

**Supplementary Fig. 5: Survival of the *kcng4b* mutant is affected after 4-aminopyridine (4-AP) treatment.**

To test the toxicity of 4-AP, embryos were treated with different concentrations of 4-AP from the 2-8 cell stage onward. Data are represented as Mean ± SD. N=3; total number of embryos 36/genotype/treatment group. A p-value of ≤ 0.05 is considered significant. A) **Survival rate for 1 mM 4-AP:** On day 1 of treatment, the survival rate of *kcng4b^-/-^* embryos was slightly lower than that of both WT and *kcnb1^-/-^* embryos. However, no difference in survival rates was observed between the WT and Kv2.1 mutants on day 2. In contrast, all *kcng4b* mutant embryos had died by day 3. Statistics were calculated using a mixed-effect model followed by Dunnett’s multiple comparison test. B) **Survival rate for 2 mM 4-AP:** The survival of *kcng4b* mutants was significantly compromised when treated with 2 mM 4-AP. At day 1, *kcng4b* mutants showed a survival rate of less than 60%, compared to the 80% survival rate of WT and *kcnb1^-/-^* embryos. By day 2, all kcng4b*^-/-^* embryos died, but no changes were observed in the survival rates of WT and kcnb1*^-/-^* embryos on days 2 and 3. Statistics were calculated using a mixed-effect model followed by Dunnett’s multiple comparison test. C) **Survival rate for 4 mM 4-AP:** Treatment with 4 mM 4-AP was detrimental to all three genotypes (WT, *kcng4b^-/-^,* and *kcnb1^-/-^*). The *kcng4b^-/-^* embryos were dead after 1 day of treatment, while the survival rate of WT and *kcnb1^-/-^* was between 50-60%. However, both WT and kcnb1*^-/-^* embryos died on day 2. Statistics were calculated using a non-parametric Kruskal-Wallis test followed by Dunn’s multiple comparison test.
