## Supplementary Table 1 for "From patterning to secretion: Kv2.1 subunits as regulators of zebrafish hatching gland morphogenesis and function"

| Gene | Expression level ratio,  *kcnb1^-/-^*/WT, SD  28 hpf | p-value | Expression level ratio,  *kcnb1^-/-^*/WT, SD  50 hpf | p-value | Expression level ratio,  *kcnb1^-/-^*/WT, SD  76 hpf | p-value |
| --- | --- | --- | --- | --- | --- | --- |
| *ctslb* | 1.165 ±0.204 | 0.329 | 1.010 ±0.379 | 0.968 | **1.99 ±0.374** | **0.0256** |
| ***he1.1*** | **0.645 ±0.042** | **0.0049** | **0.506 ±0.145** | **0.0439** | **4.959 ±0.353** | **0.0286** |
| *cd63* | 0.959 ±0.224 | 0.7754 | **1.173 ±0.045** | **0.0035** | 1.100 ±0.255 | 0.4911 |
| ***klf17*** | **0.265 ±0.028** | **0.0021** | 1.078 ±0.089 | ns | **0.681 ±0.078** | **0.0132** |
| ***foxe3*** | **0.101 ±0.031** | **0.0046** | **0.489 ±0.051** | **0.0057** | 0.625±0.340 | 0.4041 |
| ***pitx2*** | **0.297 ±0.23** | **0.0269** | **0.596 ±0.159** | **0.04** | **5.656 ±0.595** | **0.0014** |
| *xbp1* | 1.091 ±0.025 | 0.4744 | **1.154 ±0.066** | **0.0258** | 1.256 ±0.083 | 0.2613 |
| *bhlha15* | 1.474 ±0.197 | 0.0841 | 1.202 ±0.688 | 0.6414 | 1.468±0.719 | 0.3548 |
| *cdh1* | 0.983 ±0.233 | 0.8977 | 1.174 ±0.117 | 0.0836 | 1.286 ±0.122 | 0.1395 |
| *spint2* | 0.810 ±0.166 | 0.1229 | 1.059 ±0.257 | 0.6915 | **1.334 ±0.023** | **0.0059** |
| *zp2* | 2.587±0.347 | 0.0147 | - | - | - | - |

| Gene | Expression level ratio,  *kcng4b^-/-^*/WT, SD  28 hpf | p-value | Expression level ratio,  *kcng4b^-/-^*/WT, SD  50 hpf | p-value | Expression level ratio,  *kcng4b^-/-^*/WT, SD  76 hpf | p-value |
| --- | --- | --- | --- | --- | --- | --- |
| *ctslb* | 0.779 ±0.218 | 0.4084 | 0.820 ±0.091 | 0.3900 | 1.228 ±0.135 | 0.0843 |
| *he1.1* | 0.676 ±0.315 | 0.2284 | 0.800 ±0.076 | 0.1123 | **1.608 ±0.001** | **0.0049** |
| *cd63* | 1.088 ±0.235 | 0.5205 | **0.757 ±0.021** | **0.0063** | **1.404 ±0.296** | **0.0177** |
| *klf17* | 0.941 ±0.237 | 0.6925 | 1.855 ±0.150 | 0.1125 | 1.057 ±0.081 | 0.7394 |
| *foxe3* | 1.094 ±0.216 | 0.7375 | 2.013 ±0.110 | 0.0713 | **2.235±0.603** | **0.0305** |
| *pitx2* | **0.845 ±0.047** | **0.0369** | 2.230 ±0.054 | 0.1551 | **1.630 ±0.202** | **0.0307** |
| *xbp1* | 1.084 ±0.218 | 0.4526 | 1.114 ±0.042 | 0.1627 | 1.040 ±0.045 | 0.5606 |
| *bhlha15* | 0.837 ±0.151 | 0.1808 | 0.639 ±0.206 | 0.1662 | 2.073±0.179 | 0.1557 |
| *cdh1* | **0.581 ±0.075** | **0.0140** | 0.997 ±0.316 | 0.9861 | **0.687 ±0.142** | **0.0277** |
| *spint2* | 1.104 ±0.137 | 0.5444 | 0.949 ±0.023 | 0.2669 | 0.977 ±0.188 | 0.8975 |
| *zp2* | 2.858±0.539 | 0.0133 | - | - | - | - |
